# The coupling between healthspan and lifespan in *Caenorhabditis* depends on complex interactions between compound intervention and genetic background

**DOI:** 10.1101/2022.01.15.476462

**Authors:** Stephen A. Banse, E. Grace Jackson, Christine A. Sedore, Brian Onken, David Hall, Anna Coleman-Hulbert, Phu Huynh, Theo Garrett, Erik Johnson, Girish Harinath, Delaney Inman, Suzhen Guo, Mackenzie Morshead, Jian Xue, Ron Falkowski, Esteban Chen, Christopher Herrera, Allie J Kirsch, Viviana I. Perez, Max Guo, Gordon J. Lithgow, Monica Driscoll, Patrick C. Phillips

**Author notes:** Equal author contributions (Stephen A. Banse, E. Grace Jackson, Christine A. Sedore). Corresponding authors’ e-mail addresses.

## Abstract

Aging is characterized by declining health that results in decreased neuromuscular function and cellular resilience. The relationship between lifespan and health, and the influence of genetic background on that relationship, has important implications in the development of anti-aging interventions. Here we combined survival under thermal and oxidative stress with swimming performance, to evaluate health effects across a nematode genetic diversity panel for three compounds previously studied in the *Caenorhabditis* Intervention Testing Program – NP1, propyl gallate, and resveratrol. We show that oxidative stress resistance and thermotolerance vary with compound intervention, genetic background, and age. The effects of tested compounds on swimming locomotion, in contrast, are largely species-specific. Additionally, thermotolerance, but not oxidative stress or swimming ability, correlates with lifespan. Our results demonstrate the importance of assessing health and lifespan across genetic backgrounds in the effort to identify reproducible aging interventions.

## Introduction

Age-associated health deterioration results in increased prevalence of numerous diseases^1^, reduced muscle function^2^, movement^3^, and resilience to both internal and external stressors^4^. Aging is also a principal risk factor for mortality, with hazard rates increasing throughout adulthood. However, the underlying functional relationship between health and lifespan, and in particular how this relationship is modulated by genetic background, is still somewhat surprisingly unclear. One challenge in dissecting this relationship is the difficulty in defining health in model systems. For human health, integrative approaches using physical, cognitive, or physiological performance are used as a proxy for health state. These approaches include the Short Physical Performance Battery (an assessment of gait speed, chair stand, and balance in the elderly^5–7^), Frailty Index (assessed by presence of disease, physical disability, and cognitive decline^8,9^), and the Healthy Aging^10^, Successful Aging^11^, and Cognitive Frailty indices^12^. Parallel approaches have been suggested for invertebrate models (e.g., *Drosophila* locomotion^13^ vs human treadmill testing) where genetics can be controlled, thus facilitating study of both the relationship between health and lifespan, and the influence of genetic background.

*Caenorhabditis elegans* is a widely used research model that experiences age-dependent declines in a variety of physiological processes^14,15^. A number of health assays have been employed in *C. elegans* to quantify functional declines. Among the most commonly used health-related phenotypes are stress resistance and body movement^14–21^ (measures that broadly align with health and intrinsic capacity^12,22^), pharyngeal pumping^23,24^, and lipofuscin accumulation^21,25^. These measures have been used to probe the relationship between health and lifespan^19,26,27^. For example, measures of physiological function like thermotolerance and oxidative stress resistance positively correlate with longevity^19,28,29^. Motility, a measure of neuromuscular function, is positively correlated with lifespan in some but not all studies^3,15,16^.

While lifespan and the maintenance of health appear correlated, whether they are causally linked is hotly debated. There are a number of definitions of healthspan, but they are generally defined as the length of time before a precipitous loss in stress resistance and physiological function. Previous studies have shown that compound interventions that improve healthspan do not necessarily impact lifespan^30–32^. Recent work that used a definition of healthspan based on population maximum lifespans suggests that in a number of long-lived mutants, including well-characterized IIS/IGF signaling pathway mutants, healthspan can be largely uncoupled from lifespan^17^. Subsequent work and reanalyses have contradicted these results, however, finding that healthspan, at least in long-lived *daf-2* mutants, is maintained proportionally with lifespan, even as other long-lived mutants exhibit attenuated health outcomes^16^. On top of these complications, the influence of genetic background remains a relatively unexplored variable because studies have largely been constrained to a single isogenic line, namely *C. elegans* laboratory strain N2. However, genetic background plays a critical role in determining the effects of different pharmacological interventions on lifespan^33^, and compounds may affect health in species- and even population-specific ways.

The *Caenorhabditis* Intervention Testing Program (CITP) is a collaborative approach to addressing the complexities inherent in testing pharmaceutical impact on longevity and health across genetic backgrounds by exploiting the diversity of the *Caenorhabditis* genus. Using three *Caenorhabditis* species that represent genetic variation similar to that between mice and humans^34–36^, the CITP implements identical protocols at three independent sites to screen for small molecule lifespan and healthspan effects, with the goal of minimizing lab-to-lab variability to generate high quality, reproducible results. Our published data demonstrate the efficacy of our approach: in an initial screen of 12 compounds for longevity effects, we identified 6 that reproducibly promote lifespan in at least one of the tested species^33^.

Here we set out to apply CITP protocols to answer three questions for compounds with potential anti-aging effects: (1) When a compound extends lifespan, does it do so by fundamentally slowing the aging process, resulting in broad health benefits? (2) Do compound interventions promote health benefits in genetic backgrounds that do not exhibit lifespan extension? and (3) What is the relationship between health measures and lifespan, and which health measures are most reproducible and informative for future CITP compound evaluation? To answer these questions we determined health effects for three pro-longevity compounds from our previous screens^33,37^:: NP1, propyl gallate, and resveratrol. We assessed health by measuring survival under heat stress, survival under oxidative stress, and swimming performance. We find that swimming ability and oxidative stress resistance are highly reproducible across labs, but that compound intervention effects on oxidative stress and swimming ability do not correlate with lifespan effects across genetic backgrounds. In contrast, compound interventions do intersect with thermotolerance in a manner that correlates with longevity. Our results demonstrate the value of assessing health declines across genetically diverse test sets in the search for reproducible anti-aging interventions.

### Independent and combined effects on health and aging

Aging is marked by a progressive decline in physiological function and an increase in hazard rate^38^. An anti-aging intervention treating the root cause(s) of aging would slow those progressive changes, resulting in larger relative improvements late in life, rather than simply being stimulatory at all ages (see Figure 1). Anti-aging interventions therefore alter the slope of a measure over time and are detectible statistically as an age-by-compound interaction. We prefer this slope-based approach to traditional healthspan metrics because it removes hits that are merely general stimulants. Anti-aging compounds may have different effects on lifespan and health if: (1) they are separable phenotypes, (2) there are multiple causes of aging, or (3) the intervention acts on a symptom of aging instead of on the underlying cause. A compound’s effects on health and aging can therefore be classified along two axes and into four different categories (Figure 1). The first category (upper right-hand quadrant), represents both lifespan extension and a reduced rate of health decline, resulting in a healthy lifespan extension. The second category (lower right-hand quadrant) represents health preservation at the cost of lifespan. The third category (lower left-hand quadrant) represents general toxicity, with a decrease in lifespan and an increase in the rate of health decline. Interventions that fall into the final category extend lifespan while accelerating health decline (upper left-hand quadrant), which results in a relative extension of the gerospan^17^. Evaluation of data within this conceptual framework enables us to evaluate a compound’s effect on both lifespan and multiple healthspan parameters as measured across a variety of diverse genetic backgrounds.

**Figure 1.**
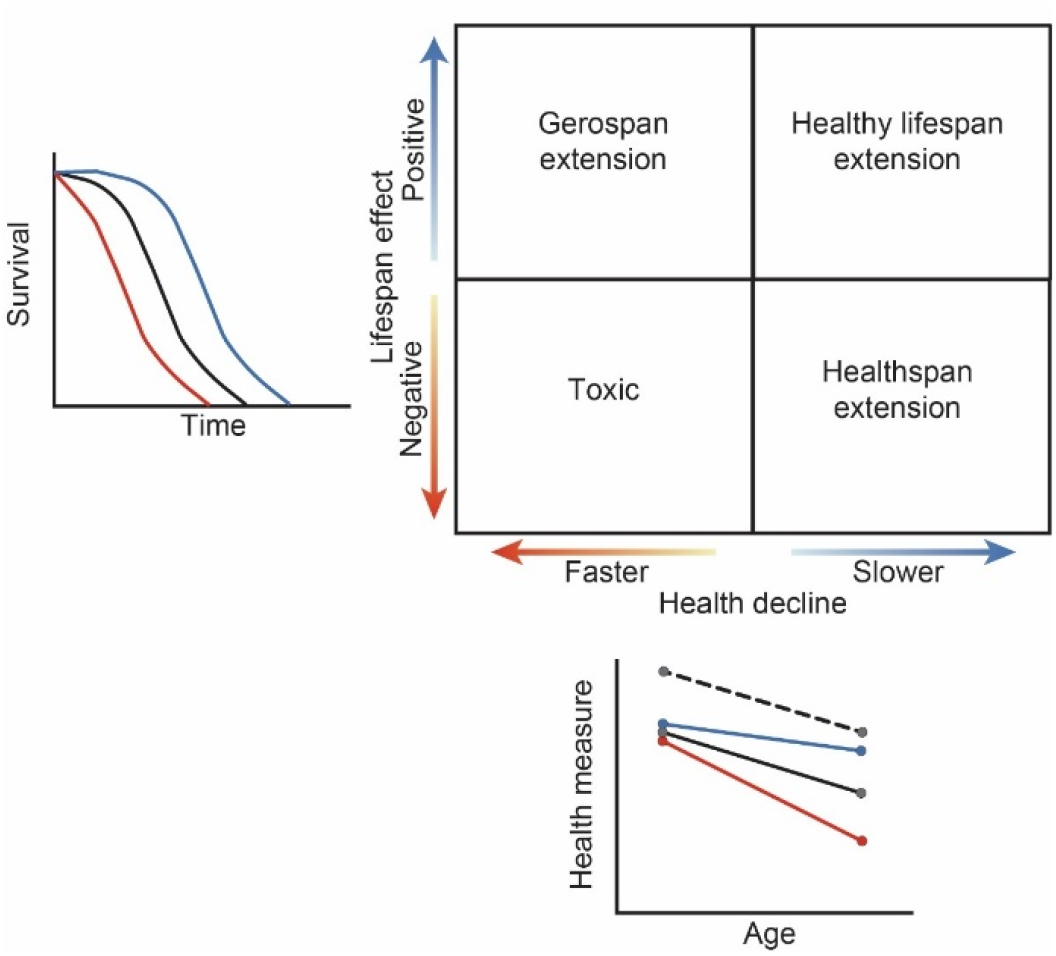
Compound effects on lifespan and health: A qualitative diagram of possible outcomes for compound effects on lifespan and on healthspan. Lifespan is represented as a change in median survival, while healthspan is represented in the relative rate of decline as compared to the control. The black lines show control, while the blue lines depict slowed aging, and the red lines depict accelerated aging. For health measures, the black dashed line shows the effects of an intervention that is generally stimulatory but does not alter the aging process. Depending on the effect size and direction, each healthspan, compound, and strain combination will fall into one quadrant: lifespan and healthspan extending, healthspan extending, gerospan extending, or toxic. The solid lines between the quadrants indicates no change from the control for a given measure.

## Results

To evaluate compound effects in this framework we selected three compounds (NP1, resveratrol, and propyl gallate) previously shown by the CITP to have positive effects on lifespan in *C. elegans*, although not in *C. briggsae*^33^. NP1 is believed to prolong lifespan by acting as a dietary restriction mimetic^39^. Similarly, resveratrol may also extend lifespan via a dietary restriction mechanism^40^, although the mode of induction for resveratrol is probably different than for NP1. Propyl gallate likely acts through a wholly distinct mechanism and was initially tested for lifespan effects in *C. elegans* based on its antioxidant properties^41^.

### Oxidative stress response varies in a compound-, strain-, and agespecific manner

Cells experience oxidative stress due to metabolic activity and environmental stressors. The ability to resist oxidative stress is fundamental to the maintenance of cellular function^42^, and is therefore a useful measure of health via stress resistance. We sought to assess whether compounds that showed species- and strain-specific effects on lifespan would exhibit similar patterns of effects on oxidative stress resistance, or if compounds could potentially confer health effects in strains that did not show increases in lifespan. In particular, we tested oxidative stress resistance in the presence of paraquat at two ages: early mid-life (adult day 6 for *C. elegans*, adult day 8 for *C. briggsae*; time points selected based on our previously detailed survival analyses in these diverse backgrounds) and late mid-life (adult day 12 for *C. elegans*, adult day 16 for *C. briggsae*) to investigate whether compound effects on oxidative stress resistance were age-dependent. Specifically, we wanted to assess if longevity-promoting compounds could ameliorate age-related declines in oxidative stress resistance, or if they merely increased oxidative stress resistance overall. Later ages were tested for *C. briggsae* strains because they display a later age of senescent decline in general.

We found a minimal effect of DR mimetic NP1 on oxidative stress resistance in all tested strains across both species (Figure 2a; Supplemental Figure 2). In particular, the strong species-specific effect that we previously reported for lifespan with NP1 treatment^33^ was not evident for oxidative stress resistance. More specifically, one strain, *C. briggsae* AF16, showed a small age-related increase in oxidative stress resistance (no significant effect at early mid-life; 15% increase in median survival at late mid-life, *p*=0.0288). A second strain, *C. elegans* MY16, demonstrated an age-by-compound interaction, with a moderately negative oxidative stress response at early mid-life (25% decrease in median survival, *p*=0.00135), but a robust positive response at late mid-life age (61% increase in median survival, *p*=0.00477). *C. briggsae* HK104 had an overall robust decrease in oxidative stress resistance at both ages (33-35% decrease in median survival, *p*<0.001). The remaining three strains, *C. elegans* N2 and JU775, and *C. briggsae* JU1348, showed no significant effect on oxidative stress resistance following NP1 treatment. We conclude that NP1 treatment in genetically diverse backgrounds induces a range of somewhat variable oxidative stress resistance responses. Importantly, for NP1 oxidative stress response does not correlate well with previously reported longevity benefits across strains.

**Figure 2.**
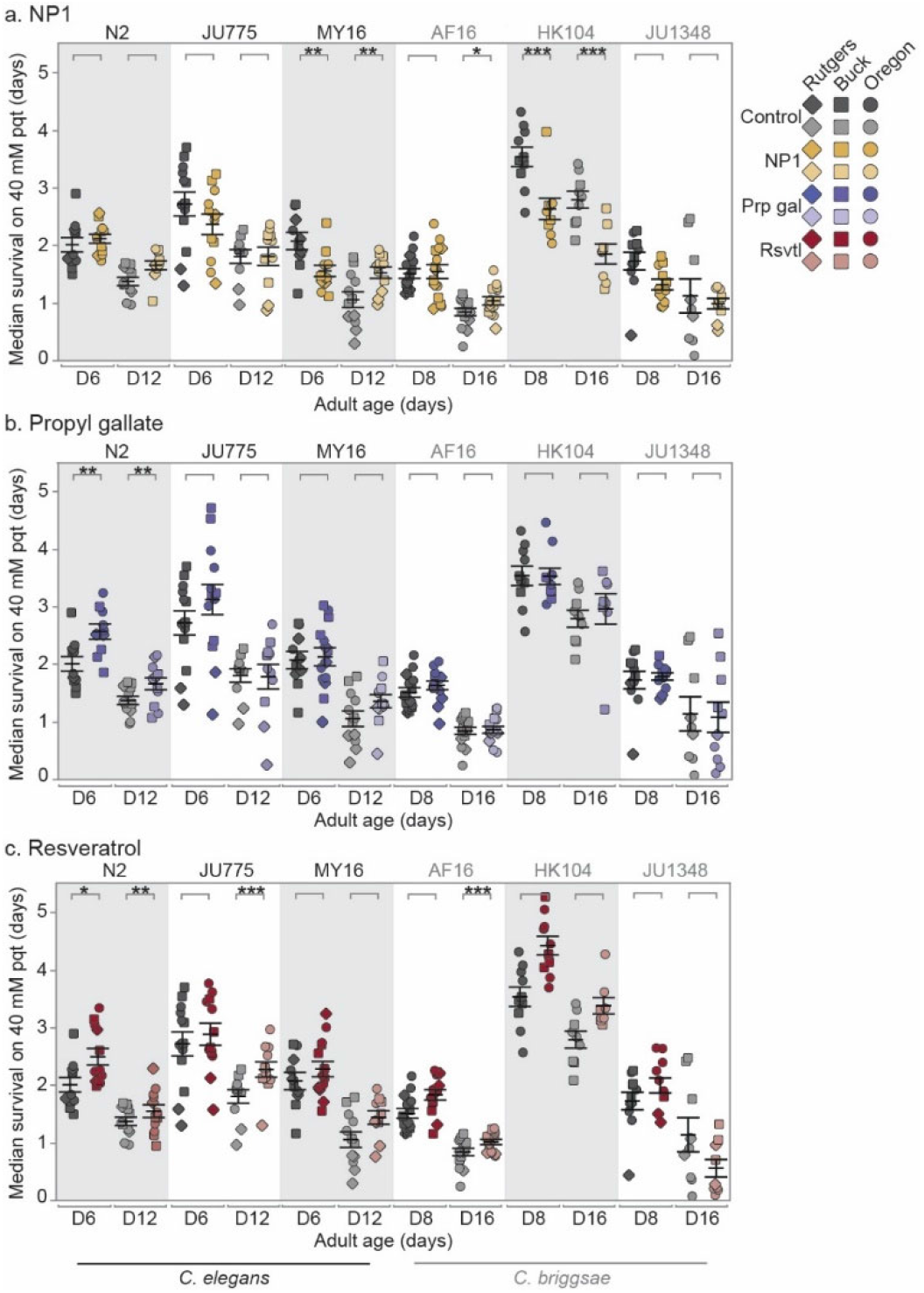
Compound effects on oxidative stress resistance: The effect of adult exposure to (a) NP1, (b) propyl gallate, and (c) resveratrol on median survival under oxidative stress conditions, beginning at day 6 and 12 (*C. elegans*), or day 8 and 16 (*C. briggsae*) of adulthood. Three strains were tested from each species: *C. elegans* strains N2, MY16, and JU775 (black text), and *C. briggsae* AF16, ED3092, and HK104 (gray text). Each point represents the median survival on 40 mM paraquat of an individual trial plate, control (vehicle only - gray) or compound treated (color). The bars represent the mean +/- the standard error of the mean. Replicates were completed at the three CITP testing sites (square – Buck Institute, circle – Oregon, and diamond – Rutgers). Asterisks represent *p*-values from the CPH model such that \*\*\*\**p*<0.0001, \*\*\**p*<0.001, \*\**p*<0.01, and \**p*<0.05.

Our previously published lifespan results showed that the antioxidant propyl gallate had a weak but positive effect on lifespan in *C. elegans*, and no effect on lifespan in *C. briggsae*^33^. Much like NP1, we found that in most cases propyl gallate did not lead to increased oxidative stress resistance (five of the six strains tested). Only one strain, *C. elegans* N2, showed an overall increase in oxidative stress resistance, which was more robust in early midlife (36% increase in median survival at early mid-life, *p*=0.00128; 24% increase at late mid-life, *p*=0.00323).

Finally, we tested the effect of resveratrol on oxidative stress resistance, another compound that we showed confers a species-specific lifespan response^33^. Unlike NP1 and propyl gallate, resveratrol had a widespread effect, and significantly increased oxidative stress resistance in three of the six strains tested (Figure 2c). *C. elegans* N2 had an increase in oxidative stress resistance at both ages (27% increase in median survival at early mid-life, *p*=0.01061; 17% increase at late mid-life, *p*=0.00688), while survival increased in *C. elegans* JU775 in an age-dependent manner (no significant effect at early mid-life; 13% increase in median survival at late mid-life, *p*=0.001988). *C. elegans* MY16, along with *C. briggsae* AF16 and HK104 trended towards a general increase in oxidative stress resistance, although not all effects were significant. The remaining strain, *C. briggsae* JU1348, showed no significant change in oxidative stress resistance with resveratrol treatment.

Overall, compounds previously observed to increase lifespan have little to no effect on healthspan as measured by oxidative stress resistance, exept for resveratrol which did so in a species specific manner.

### Thermotolerance varies in a compound-, species-, strain-, and age-specific manner

Thermotolerance has been implicated as an important correlate of increased longevity^19,43^. To determine the thermotolerance effects of the test set of compounds, we aged animals at 20°C in the presence of the compound and then subsequently measured survival after shifting them to 32°C on compound-free plates (see Supplemental Figure 3 for Kaplan-Meier survival curves).

We found that NP1 ameliorated age-related thermotolerance decline in two of the six strains tested (Figure 3a), *C. elegans* N2 and MY16 (no significant change at early mid-life; 42-53% increase in median survival at late mid-life; *p* =0.00611-N2; *p*<0.0001 - MY16). Conversely, we found that *C. briggsae* HK104 exhibited accelerated loss of thermotolerance late in life (no effect at early mid-life; 28% decrease in median survival at late mid-life, *p*<0.001).

**Figure 3.**
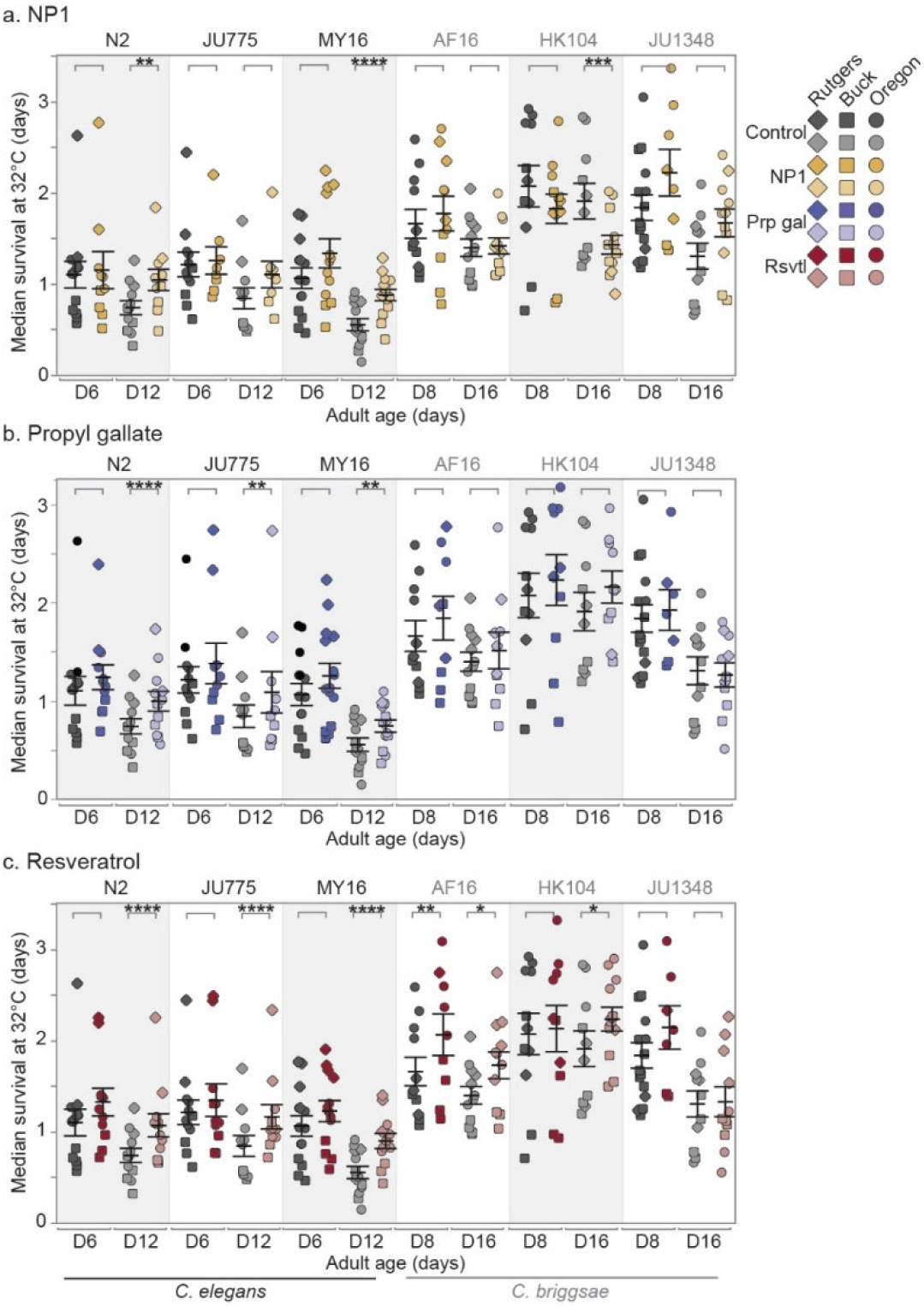
Compound effects on thermotolerance: The effect of adult exposure to (a) NP1, (b) propyl gallate, or (c) resveratrol on thermotolerance, specifically median survival at 32°C. Thermotolerance assays were run beginning on day 6 and 12 (*C. elegans*) or day 8 and 16 (*C. briggsae*) of adulthood. Three strains were tested from each species: *C. elegans* strains N2, MY16, and JU775 (black text), and *C. briggsae* AF16, ED3092, and HK104 (gray text). Each point represents the median survival at 32°C of an individual trial plate, either control (vehicle only - gray), or compound treated (color). The bars represent the mean +/- the standard error of the mean. Replicates were completed at the three CITP testing sites (square – Buck Institute, circle – Oregon, and diamond – Rutgers). Asterisks represent *p*-values from the CPH model such that \*\*\*\**p*<0.0001, \*\*\**p*<0.001, \*\**p*<0.01, and \**p*<0.05.

Treatment with propyl gallate had a moderate positive late life increase in thermotolerance in all three *C. elegans* strains tested (Figure 3b; no significant effect at early mid-life; 9-34% increase in median survival at late mid-life, *p*<0.0001 - N2, *p*=0.00138 - JU775, *p*=0.00238 - MY16). Within this group, the effect size varied in a strain-specific matter. In contrast, thermotolerance in the *C. briggsae* strains tested was not affected by propyl gallate treatment.

Finally, resveratrol caused a significant increase in thermotolerance in both species and five of the six strains (Figure 3c). More precisely, we saw a compound-by-age interaction in four of the six strains tested. These strains exhibited an increase in ability to withstand thermal stress in an age-dependent manner (no effect at early mid-life; 12-51% increase in median survival at late mid-life, *p*<0.0001 for all *C. elegans* strains, *p*<0.05 for *C.briggsae* HK104). *C. briggsae* AF16 was positively affected overall by resveratrol treatment, with an increase in survival at both ages tested (34% increase in median survival at early midlife, *p*=0.00186; 17% increase at late mid-life, *p*=0.03046), although the size of the effect decreased with age. The remaining strain, *C. briggsae* JU1348, showed no change in thermotolerance with resveratrol treatment.

Overall, for compound impact on thermotolerance in a genetically diverse test set, we find the heat resistance associated with lifespan-extending compounds varies by compound. Thermotolerance after NP1 treatment was strain- and age-dependent, while propyl gallate caused a general species- and agespecific response. Resveratrol had the most robust and widespread effect on thermotolerance, increasing survival in nearly every strain, with most exhibiting age-dependent effects, and the effect at the younger ages being nonsignificant but trending towards an increase. Notably, thermotolerance outcomes do not precisely correlate with oxidative stress results. We conclude that distinct measures of stress response can be differentially regulated by the interventions we tested.

### The effect of compounds on locomotory health is largely speciesspecific

Previous work has shown that motility decreases in aging *C. elegans*^3,15,20,44^. As locomotion is similar to assays used in human clinical health assessments, we investigated whether lifespan-extending compounds would improve motility, particularly later in life. To measure locomotion, we used the CeleST platform^44,45^ to acquire eight measures of swimming ability at days 5, 9, and 12 of adulthood, respectively (see Supplemental Figure 5 for results with the eight measures of movement). While the eight measures provide a broad range of information on the swimming of each of the strains, it was not clear which measures best capture the decline of locomotion with age across our genetic diversity panel. We therefore combined the information from all eight measures for each strain into a single multivariate composite measure using a linear discriminate analysis to weight, and combine, the individual measurements (Materials and Methods, Supplemental file 1, and online material^46^ provide an in depth description of methodology). This approach accounts for interdependency among measures and the unique movement properties of the different strains. We used a strain-specific composite score because strain differences in swimming behavior do not necessarily relate to aging per se.

With regard to locomotion, we found that NP1 had the most robust and widespread effect on swimming ability, improving locomotion in all six strains tested (Figure 4). In *C. elegans*, the effect was age-dependent, with NP1 treatment slowing the rate of decline in swimming ability in all strains. We observed the largest effect in *C. elegans* N2 (17% increase in mean swimming score at day 12, *p*<0.0001), while *C. elegans* JU775 and MY16 exhibited small but significant improvements in locomotion (2-5% increase with age, *p*<0.05 - JU775; 3% increase at adult day 9, *p*=0.0085 - MY16). Interestingly, the effect of NP1 on swimming ability in *C. briggsae* was strikingly robust. JU1348 showed a reduction in age-related locomotory decline (12-16% increase in mean swimming score at adult days 9 and 12, *p*<0.0001), while both AF16 and HK104 had an overall increase in swimming ability at all ages (9-14% increase in mean swimming score, *p*<0.0001 for all ages with the exception of HK104 day 9, where *p*=0.0028). Although NP1 conferred an overall increase in the locomotory ability of HK104, the effect was greatest in young animals, meaning that NP1 increased the relative rate of decline in strain HK104.

**Figure 4.**
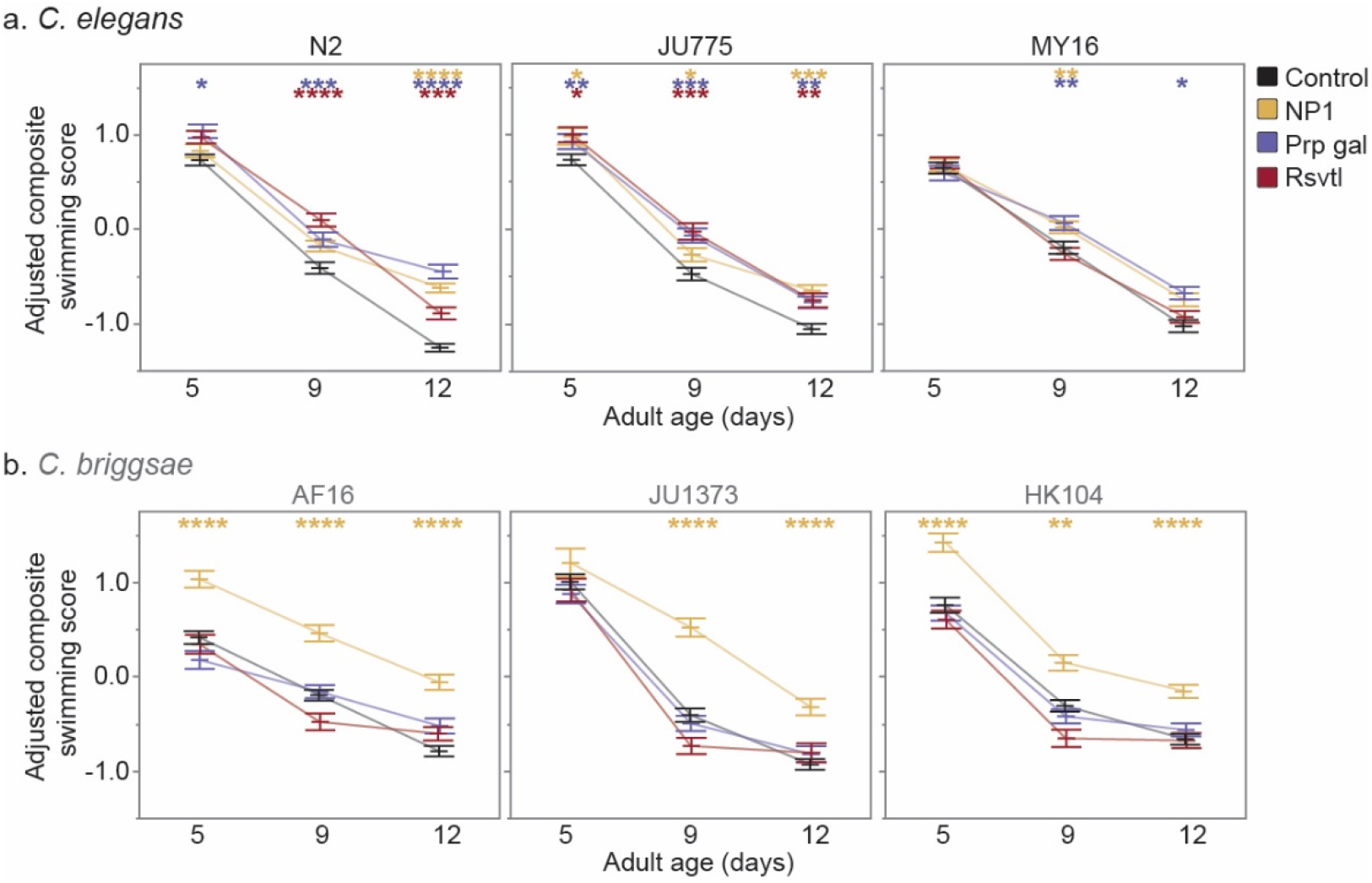
Compound effects on CeleST composite scores: The effect of adult exposure to NP1, propyl gallate, or resveratrol on overall swimming ability with age in three *C. elegans* strains (N2, JU775, MY16), and three *C. briggsae* strains (AF16, JU1348, HK104).Swimming assays were run on days 5, 9, and 12 of adulthood. Bars represent the mean +/- the standard error of the mean. Adjusted swimming score values were normalized to the strain mean value. Two biological replicates were completed at each of the three CITP testing sites. Asterisks represent *p*-values from the linear mixed model such that \*\*\*\**p*<0.0001, \*\*\**p*<0.001, \*\**p*<0.01, and \**p*<0.05.

We found that propyl gallate treatment improved swimming ability in a species-specific manner, similar to the propyl gallate effect on lifespan (Figure 4). All three *C. elegans* strains showed an age-related improvement in swimming ability, with the effect size increasing at the older ages tested. Propyl gallate improved locomotory ability in N2 by 5% (*p*=0.0105) at adult day 5, 7% (*p*=0.0005) at day 9, and 21% (*p*<0.0001) at day 12. This trend was also observed in both JU775 and MY16 to a lesser extent (4-6% maximum increase in mean composite swimming score during adult days 9 and 12; *p*≤0.0003 - JU775, *p*<0.05 - MY16), indicating that propyl gallate reduces the rate of age-related locomotory decline in *C. elegans*. In contrast, propyl gallate treatment had no effect on swimming ability in any *C. briggsae* strain tested.

We found the effect of resveratrol on locomotory ability was also species-specific (Figure 4). Specifically, resveratrol treatment improved swimming ability with age in *C. elegans* JU775 and N2 (5-11% increase in mean composite swimming score at day 9, 4-10% increase at day 12; *p*≤0.0001 - N2, *p*≤0.0015 - JU775), slowing the age-related decline in locomotion in both strains. This effect, however, is dependent on genetic background, as swimming ability in *C. elegans* MY16 was not affected by resveratrol. Likewise, resveratrol treatment had no effect on swimming in any *C. briggsae* strain.

Overall, we found that compounds that extend lifespan in particular strains also improve locomotion when administered to those same strains. It is noteworthy, however, that we also found that NP1, which did not have a lifespan effect in the *C. briggsae* genetic background, could nonetheless enhance adult swimming ability. Thus, NP1 extends locomotory health more broadly than it enhances longevity in a genetically diverse test set.

### Thermotolerance, but not oxidative stress or swimming ability, correlates with lifespan

Do health measurements correlate with lifespan, especially in response to a pharmacological intervention? Because our health assays were terminal, we do not have health and lifespan measurements for the same individuals. However, we were able to correlate each health measure score on a combined strain, compound, and age basis (Supplemental Figure 6) and found that thermotolerance was the most predictive of median lifespan, particularly at days 12 (*C. elegans)* and 16 (*C. briggsae*) of adulthood (R^2^=0.58 at early mid-life, R^2^=0.83 at late mid-life). In contrast, both oxidative stress resistance and swimming ability did not correlate well with median lifespan, with swimming ability being a particularly poor predictor of the effect of a compound on longevity (R^2^=0.08, 0.097, 0.048, for days 5, 9, and 12 of adulthood, respectively). Our data suggest that, despite experimental variability in thermotolerance outcomes, thermal resistance might be the best proxy for longevity promoting drugs, an idea that remains to be tested more extensively.

### Swimming ability and oxidative stress resistance, but not thermotolerance, are highly reproducible across labs

We observed some variability within our healthspan datasets and statistically evaluated the sources of this variation (Table 1). For oxidative stress resistance, ~34% of the total variance is attributed to genetic background, ~7.8% of total variance is attributed to lab-specific effects, and ~9.8% of the total variance is a result of variation within labs. This distribution of variation is similar to the variation we observed in our previously published lifespan datasets^33,37^. The sources of variation within the swimming dataset are weighted towards individual variation, with ~79.2% of total variance attributable to differences in individual swimming performance. Variability at the genetic level only accounts for 5.7% of the variance, with nearly half of that (2.4%) attributable to species differences. Furthermore, among-lab differences only contributed to ~5.8% of the total variation, and within lab variance contributed to ~9.3% of the total variation. Both oxidative stress resistance and swimming are thus reproducible across labs.

We found thermotolerance outcomes to be variable in comparison to longevity, locomotion and oxidative stress measures. For thermotolerance, ~24.4% of the total variation is attributed to genetic background, while we find ~28.5% of variation is attributable to among lab differences, and ~15.8% to variation within labs. Variability may reflect difficult-to-control environmental factors such as localized/fluctuating differences in incubator temperature (or temperature distribution), or humidity variation. Regardless of root cause, however, data reveal that even against the backdrop of variability in thermotolerance, this measure may serve as a plausible tool for assessing the chances that a given intervention might exert longevity benefit.

## Discussion

The ultimate goal of exploiting model organisms to screen for antiaging interventions is to identify treatments that might translate to healthy lifespan extension in humans (Figure 1). Few people are interested in living to be 120 years old if that means continuing health declines seen in most 90-year-olds, but most would be happy to live to 100 with a health state similar to 60-year-olds. Despite widespread application of the longevity screening approach, it remains somewhat unclear as to whether the goal of translating model organism research to healthy human aging is achievable, in part because the relationship between healthspan and lifespan varies based on how/when health is measured, analyzed, and interpreted^16,17,47,48^. Using varying measures of health, some compound interventions exhibit independent effects on health and lifespan^30–32^. Whether this disconnect between phenotypes is surprising reflects typically unspoken assumptions regarding the mode of action of interventions, and the underlying causation of the age-dependent phenotypes of health and lifespan.

**Table 1.** Comparison of reproducibility of health measurements from oxidative stress, thermotolerance, and CeleST assays within and between labs.

| Source of Variation | Oxidative Stress | Thermo | CeleST |
| --- | --- | --- | --- |
| **Genetic Variation** | **34.0** | **24.4** | **5.7** |
| Among species | 0.0 | 0.6 | 2.4 |
| Among strains w/in species | 29.2 | 3.7 | 0.9 |
| Species x age | 0.1 | 1.4 | 0.1 |
| Species x compound | 0.0 | 0.0 | 0.8 |
| Species x age x compound | 0.0 | 15.3 | 0.2 |
| Strain x age | 0.0 | 1.0 | 0.7 |
| Strain x compound | 3.8 | 1.9 | 0.1 |
| Strain x age x compound | 0.9 | 0.6 | 0.6 |
| **Reproducibility Among Labs** | **7.8** | **28.5** | **5.8** |
| Among labs | 3.5 | 13.8 | 3.3 |
| Lab x species | 0.0 | 7.5 | 0.6 |
| Lab x strain | 3.6 | 0.8 | 1.1 |
| Lab x age | 0.5 | 4.8 | 0.6 |
| Lab x compound | 0.0 | 0.7 | 0.0 |
| Lab x age x compound | 0.1 | 0.9 | 0.2 |
| **Reproducibility Within Labs** | **9.8** | **15.8** | **9.3** |
| Among trials | 4.1 | 6.8 |  |
| Among scanners w/in trials | 0 | 3.6 |  |
| Among plates w/in scanners | 5.8 | 5.4 |  |
| Among experimenters |  |  | 0.8 |
| Among trials w/in expts |  |  | 2.0 |
| Among videos w/in trials |  |  | 6.5 |
| **Individual Variation** | **48.4** | **31.3** | **79.2** |
| **Total** | **100.0** | **100.0** | **100.0** |
| Total number of observations | 20,467 | 22,213 | 15,117 |

For example, compound interventions that treat the root cause(s) of aging would be expected to confer broad age-dependent health benefits, as well as reduced mortality. Yet, it is possible to treat symptoms of aging instead of underlying causes. In those cases, if the symptom is only related to mortality, or a particular health measure, we would expect separation between lifespan and health effects. It is therefore beneficial when characterizing anti-aging interventions to ask: are there are broad health benefits indicative of a fundamentally slowed aging process? And is that aging process influenced by genetic background?

### Genetic background can profoundly affect the efficacy of a tested intervention to promote survival or health

Although many screens for longevity extension in *C. elegans* have been published^49–52^, genetic background effects remain an unexplored variable in these screens as nematode studies have been largely confined to the canonical *C. elegans* N2 lab strain. The CITP seeks to address the issue of genetic background variability in compound screening by testing a panel of *Caenorhabditis* strains that represent similar evolutionary divergence to that found between mice and humans (^34–36^). As we have previously published, genetic background is important when assessing the effects of pharmacological interventions on lifespan^33^. Here we selected three anti-aging compounds (NP1, propyl gallate, and resveratrol) for which we had previously conducted comprehensive studies of longevity^33^ and we characterized their impact on two measures of physiological resilience, oxidative stress resistance (Figure 2) and thermotolerance (Figure 3), and on swimming ability (Figure 4), a primarily neuromuscular phenotype thought to reflect general health and vigor.

To determine if these three compounds induced lifespan extension by affecting a root cause in the aging process, we determined whether they conferred broad health benefits. We observed that if a compound extended lifespan in a particular strain, that compound typically slowed the rate of decline across the healthspan measures tested, with some exceptions (Figure 5). This raises an interesting question: if a compound is modulating a core aging process, then why are the observed compound effects dependent on the genetic background? This may result from background differences in permeability, compound turnover, pathway tuning that can vary with genetic background, or may suggest that the underlying cause(s) of aging varies across genetic backgrounds, or that the compounds are affecting symptoms of aging instead of causes of aging. This separation in phenotypes suggests that interventions that do not exhibit a benefit in one age-dependent measure may show benefits when assayed using an alternative measure. We therefore sought to determine if compound interventions promote health benefits in genetic backgrounds that do not exhibit lifespan extension. Interestingly, the strains that saw no lifespan change after compound intervention had varying and sometimes robust effects on the rate of health decline, both positively and negatively. In this way, we observe that health effects can be uncoupled from lifespan, and that anti-aging effects could be seen on health regardless of lifespan effects.

**Figure 5.**
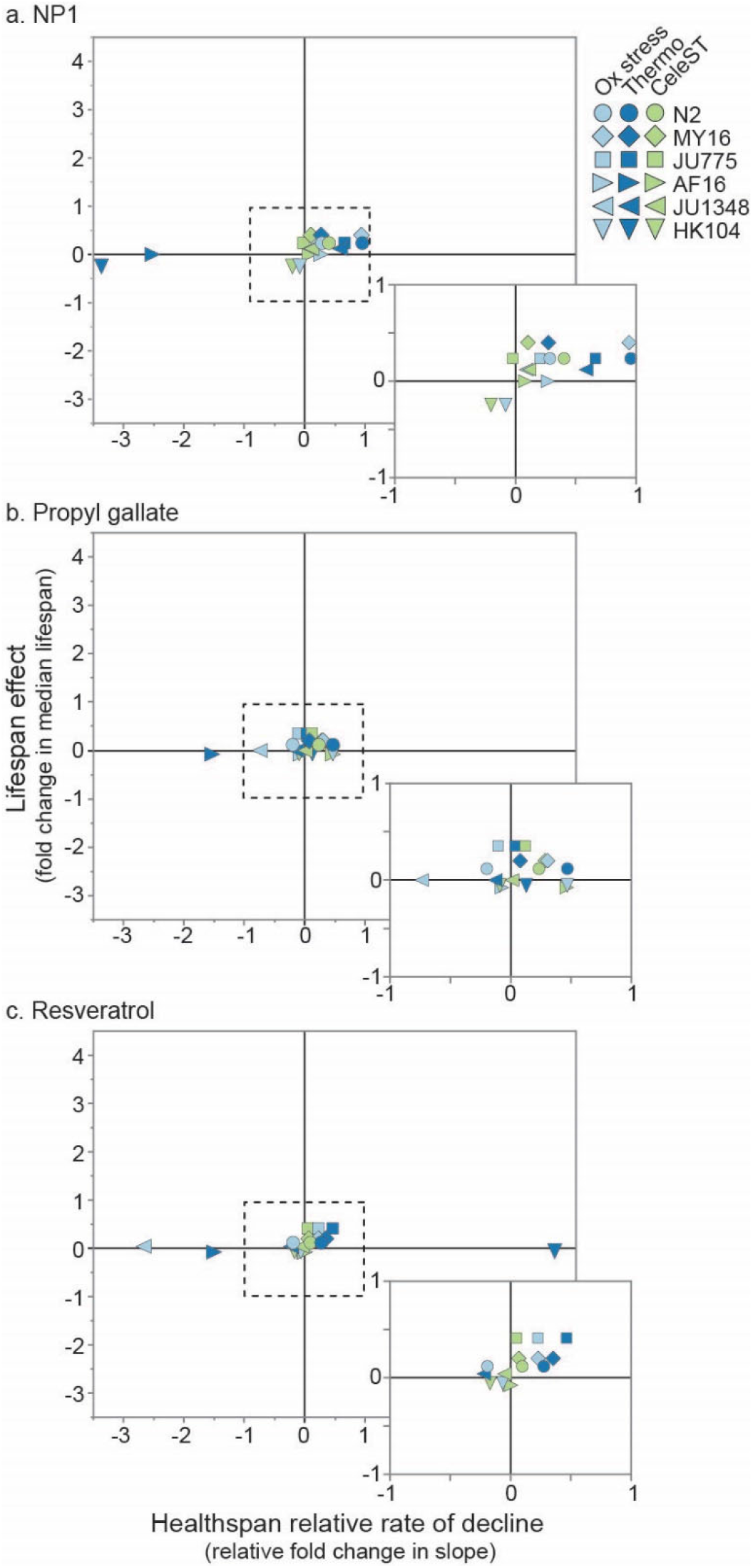
Relative compound effects on health vs. lifespan: Comparing the effect of (a) NP1, (b) propyl gallate, and (c) resveratrol on manual lifespan versus healthspan measures in two *Caenorhabditis* species. The lifespan effect is the fold change in median lifespan for a strain compared to its untreated control. For health, the relative rate of decline for each strain and compound is compared to the rate of decline for the control. Positive numbers would reflect either lifespan extension or slowed decline in the health measure, while negative numbers would reflect shortened lifespan and accelerated decline in the health measure. Each point represents a strain and health measure combination. Dotted line surrounds expanded box.

### Thermotolerance as a proxy for longevity outcomes?

Given the relationship between different health measures and lifespan, we sought to determine which health measures are most reproducible and informative for targeted screens for anti-aging treatments. In the case of the three health measures that we evaluated, thermotolerance was the only one for which compound intervention effects correlated well with lifespan effects across genetic backgrounds (Supplemental Figure 6). It is well known that increased thermotolerance corresponds to increased longevity^19,53–55^. The fact that lifespan correlates significantly with thermotolerance across compounds and genetic backgrounds suggests that the ability to maintain cellular homeostasis is a key factor in not just maintaining health, but also crucial in preventing death. Thermal stress causes unfolding of cellular proteins, and the induction of stress response pathways which induce organelle-specific molecular chaperone production^56,57^. We therefore interpret the correlation between lifespan and thermotolerance as being due to compound administration promoting protein homeostasis and therefore resulting in lifespan extension and increased thermotolerance. Despite this relationship, thermotolerance may only be used sparingly in future CITP compound screening for three reasons: (1) Given the correlation between thermotolerance and lifespan, there may be little additional benefit for using thermotolerance assays given that positive hits will likely be identified by lifespan studies alone, (2) although in principle thermotolerance provides a faster result, the approach used here actually ended up being more labor intensive than standard longevity assays, and (3) most importantly, among our three assays, thermotolerance was the least reproducible across labs (Table 1).

### Swimming ability and oxidative stress resistance reflect aging processes that are uncorrelated with lifespan

In contrast to thermotolerance, oxidative stress resistance, the other physiological resilience measure we used in these studies, does not correlate well with lifespan (Supplemental Figure 6). Even though oxidative damage has been proposed as a cause for aging^58–60^, this may not be surprising for two reasons. First, it has been shown repeatedly that antioxidant treatment does not guarantee increased lifespan, indicating that oxidative stress may not be a primary driver of mortality^30–32^, and certainly the positive and negative aspects of ROS signaling play into the outcome equation^61^. Secondly, “oxidative stress” is an umbrella term that covers a wide range of phenomena, with sub-cellular localization of the oxidative stress and differing reactive oxide species resulting in differing levels of cellular stress. Additionally, the potential for positive ROS signaling and hormesis could complicate the outcome of anti-aging interventions. In this case we only looked at a severe oxidative stressor, paraquat, and our results may have been different had we used a different oxidative stressor.

Similar to oxidative stress, we found that swimming ability, a measure of neuromuscular function^62^, also also does not correlate well with median lifespan across the different genetic backgrounds tested (Supplemental Figure 6). While oxidative stress resistance and swimming ability do not predict ultimate lifespan, these measures remain of particular interest when exploring the effect of pharmacological interventions on aging. These two measures showed high reproducibility between labs (Table 1), reflect important underlying health states, and their separability from lifespan can facilitate the identification of interventions that treat important symptoms of aging that may not be found by screening lifespan alone.

Here we have shown that pharmacological interventions result in a complex pattern of health effects, with genetic background and age both being important for overall outcomes. Our results underscore that genetic background can be a crucial determinant in the evaluation of potential anti-aging pro-resilience compounds. Relevant to human interventions, our findings emphasize that a personally tailored intervention that accounts for age, current health state, and genetic background is likely necessary for optimal efficacy. Essentially, for aging there is no medicine, just personalized medicine.

## Materials and Methods

A detailed set of standard operating procedures are available online^63^. The experimental details in brief are as follows:

### Strains

Following standard CITP protocol, the following natural isolates were obtained from the *Caenorhabditis* Genetics Center (CGC) at University of Minnesota: *C. elegans* N2, MY16, JU775; *C. briggsae* AF16, HK104, JU1348. The N2 strain used was N2-PD1073, which is a clonal line derived from the N2 strain VC2010 used to generate a new N2 reference (“VC2010-1.0)” genome^64^ that has been adopted by the CITP as a lab adapted control^37^. Worms were maintained at 20 °C on 60 mm plates with NGM and *E. coli* OP50-1 and synchronized by timed egg-lays (for full SOP see^63^). Animals were transferred to 35 mm NGM plates containing 51 μM FUdR and compound intervention (or the solvent DMSO in control plates) on the first, second and fifth day of adulthood, then once weekly when applicable, until healthspan measurements were initiated.

### Interventions

The following compounds were selected to study impacts on health based on our previous findings for nematode longevity ^33,37^: NP1 (ChemBridge), propyl gallate (Sigma-Aldrich), and resveratrol (Cambridge Chemical). The same in-plate concentrations previously tested for lifespan effects from each compound were used here: 50, 200 and 100 μM, respectively. Again, animals were exposed to compound interventions only for the duration of adulthood up until health measurements were performed.

### Selection of ages for health assays

To measure health in aging adults we selected time points that (1) showed age-dependent differences in the baseline measurements (e.g., detectable aging between the timepoints), and (2) were justifiable based on the known physiological and demographic changes in normally aging adults. For example, *C. elegans* and *C. briggsae* hermaphrodites are self-fertile, with a normal reproductive period in the absence of males lasting for the first five to eight days of adulthood^65^. We therefore used the end of the period of self-reproduction to establish an “early-mid-life” timepoint for health assays. We also selected a “late-mid-life” timepoint to correspond to approximately the 95^th^ survivorship centile ^66^ to minimize selection biases (see Supplemental Figure 1).

### Heat stress

To measure the impact of each compound on organismal tolerance of heat stress, we implemented the following augmented Lifespan Machine (LM) protocol^37,67^: 70 animals each placed on 50 mm tight-lidded petri plates with modified NGM and *E. coli* OP50-1 at 32 °C humidity for a duration of four days. For each of two biological replicates, we created two technical replicates per strain and condition (age and compound or control) per lab. *C. elegans* were tested at adult days 6 and 12 while *C. briggsae* were tested at adult days 8 and 16. A full protocol is available online^63^.

### Oxidative stress

To measure the impact of each compound on organismal resilience to exogenous oxidative stress, we implemented the following LM protocol^37,67^: 70 animals were placed onto each 50 mm tight-lidded petri plate with modified NGM containing 40 mM paraquat (or methyl viologen dichloride, from Sigma-Aldrich), 51 μM FUdR, and *E. coli* OP50-1. The duration of the paraquat-exposure assays was dependent on the starting age of the animals due to the increased rate of mortality with age. Specifically, *C. elegans* on day 6 of adulthood and *C. briggsae* on day 8 of adulthood were assayed for 16 days, while *C. elegans* at 12 days of adulthood and *C. briggsae* at 16 days of adulthood were assayed for only 7 days. For each of two biological replicates, we created two technical replicates per strain and condition (age and compound or control) per lab. A full protocol is available online^63^.

### C. elegans *Swim Test*

To measure the impact of lifespan-enhancing compounds on the age-associated decline in agility or neuromuscular function, we implemented the *C. elegans* Swim Test (CeleST; ^44^). Briefly, animals were exposed to compound intervention during adulthood as described above until CeleST measurements were collected at adult ages day 5, 9, and 12. For two biological replicates at each of the three CITP sites, 40 animals were tested per condition (age and compound or control) per strain. For full experimental protocols see our online protocol^63^. The CeleST software was used to measure eight different parameters (Wave initiation rate, Body wave number, Asymmetry, Stretch, Curling, Travel speed, Brush stroke, and Activity index) ^44,45^. To facilitate comparisons between strains and compound treatments we generated a single composite swimming score.

### CeleST Composite score

CeleST provides video-based analysis of eight separate features of locomotion, from bending rate to travel speed (Fig. 6; Ibáñez-Ventoso (2016)). However, we do not necessarily have an a priori expectation as to how each variable might change with age, or how strains may differ in age-dependent changes among the eight measures. To maximize the differences between ages across all the measurements, we used linear discriminant analysis (LDA), with the eight original measurements in each record serving as the predictor variables and age-dependent decline being the primary discriminator (Fig. 7), to reduce the number of dimensions from eight to one. Projecting the eight predictor variables onto a single axis creates the first linear discriminant function. This first linear discriminant function minimizes within-age variance, maximizes between-age variance, and maximizes the separability of the means of the ages^68^. While it cannot capture all the information provided by each original measurement, it captures as much as possible. Because it maximizes the differences between group levels (ages in our case), linear discriminant analysis is often used to predict group membership after a training dataset is used. To avoid confounding strain specific differences in swimming, our LDA was performed on the untreated control data independently for each strain to generate first linear discriminant functions for each strain. We then used the coefficients for each of the original eight predictor variables in the strain specific first linear discriminant functions as strain specific weightings of the eight measures to generate composite scores that maximized the ability to separate the animals in the control set by age. Those strain specific weightings were then used to generate a composite score of treated animals to analyze for compound and age effects. (see supplemental file 1 for in depth methodology). While the main text presents an analysis of swimming using the composite score, analyses of the individual CeleST measurements are available in the supplemental materials (Supplemental Figure 4). All relevant R-scripts and raw output are available online^46,63^.

For statistical analysis of swimming behavior, the composite score was used as the target variable of interest in mixed effects general linear models built for each strain in R using the Lme4 package (v1.1.27.1)^69^, with compound and age as fixed effects and laboratory site, research technician, experiment date, and video as nested random effects. Determination of significant age by compound interactions was made using the R car package (v3.0.11)^70^. Such an interaction where the impact size of a compound varies by age would imply an altered rate of aging for that health measure. For example, differing aging effects could be observed when we tested the healthspan effects of three of the compounds found to have positive effects on lifespan (slowed aging induced mortality) in previous CITP experiments^33^. While most of these compounds also had positive effects on overall locomotory health across the life cycle (e.g., Fig. 7), the compounds did not have dramatic age by compound effect interactions. This would suggest that although these compounds may be beneficial through general stimulatory effects on swimming, the lack of increasing locomotory heath benefits later in life suggests that the benefits are not conferred through slowed locomotory aging.

## Acknowledgements

We acknowledge all of the members of the Lithgow, Driscoll and Phillips labs for helpful discussions. We thank the CITP Advisory Committee and Ronald Kohanski (National Institute on Aging) for extensive discussion. We thank Asher Cutter, Marie-Anne Félix, and Christian Braendle for providing strains that they had directly collected. Additional strains were provided by the CGC, which is funded by NIH Office of Research Infrastructure Programs (P40 OD010440). This work was supported by funding from National Institutes of Health grants (U01 AG045844, U01 AG045864, U01 AG045829, U24 AG056052).

## Author Contributions

Conceptualization: MD, GJL, PCP, MG, SAB. Methodology: CH, EC, DH. Formal analysis: EGJ, CAS, PCP. Data curation: EGH, CAS. Funding acquisition: MG, VIP, GJL, MD, PCP. Conducted experiments: CAS, EJ, ACH, DH, BO, TG, PH, DI, MM, GH, SG, JX, AK. Visualization: CAS, SAB. Wrote the paper: SAB, CAS, EJ, ACH. Reviewed, edited, and approved the paper: All authors.

