## Supplemental Figures for "The coupling between healthspan and lifespan in *Caenorhabditis* depends on complex interactions between compound intervention and genetic background"

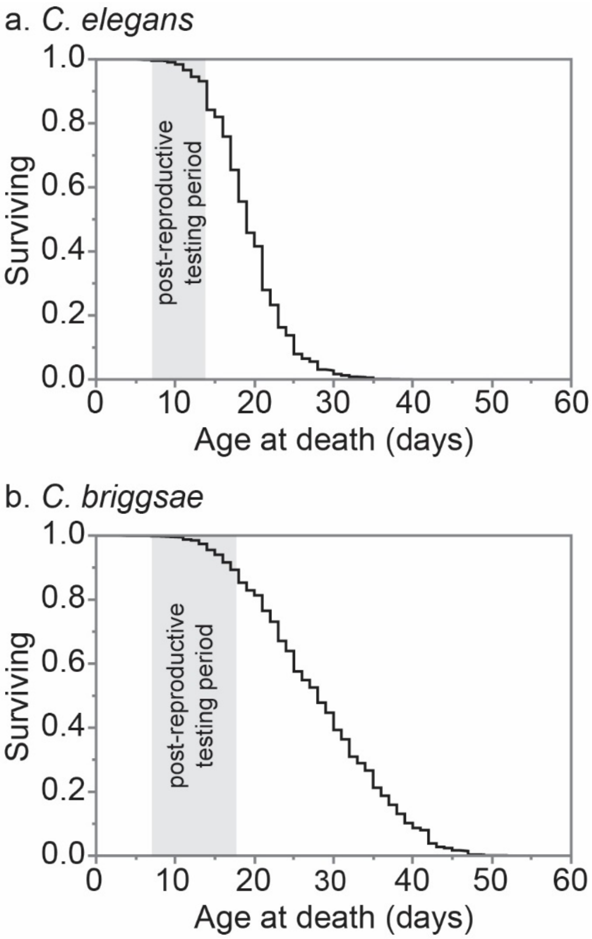


**Supplemental Figure 1. Health assay period** Kaplan-Meier lifespan curves showing overall species lifespan curves for (a) *C. elegans*, and (b) *C. briggsae*. Each curve consists of our previously published baseline survival data[33] from the three strains tested within a given species: N2, JU775, and MY16 for *C. elegans*, and AF16, JU1348, and HK104 for *C. briggsae*. Ages tested for health metrics were chosen based on the end of reproduction up until the (approximate) 5% quantile of survival under baseline conditions.

### Supplemental Figures

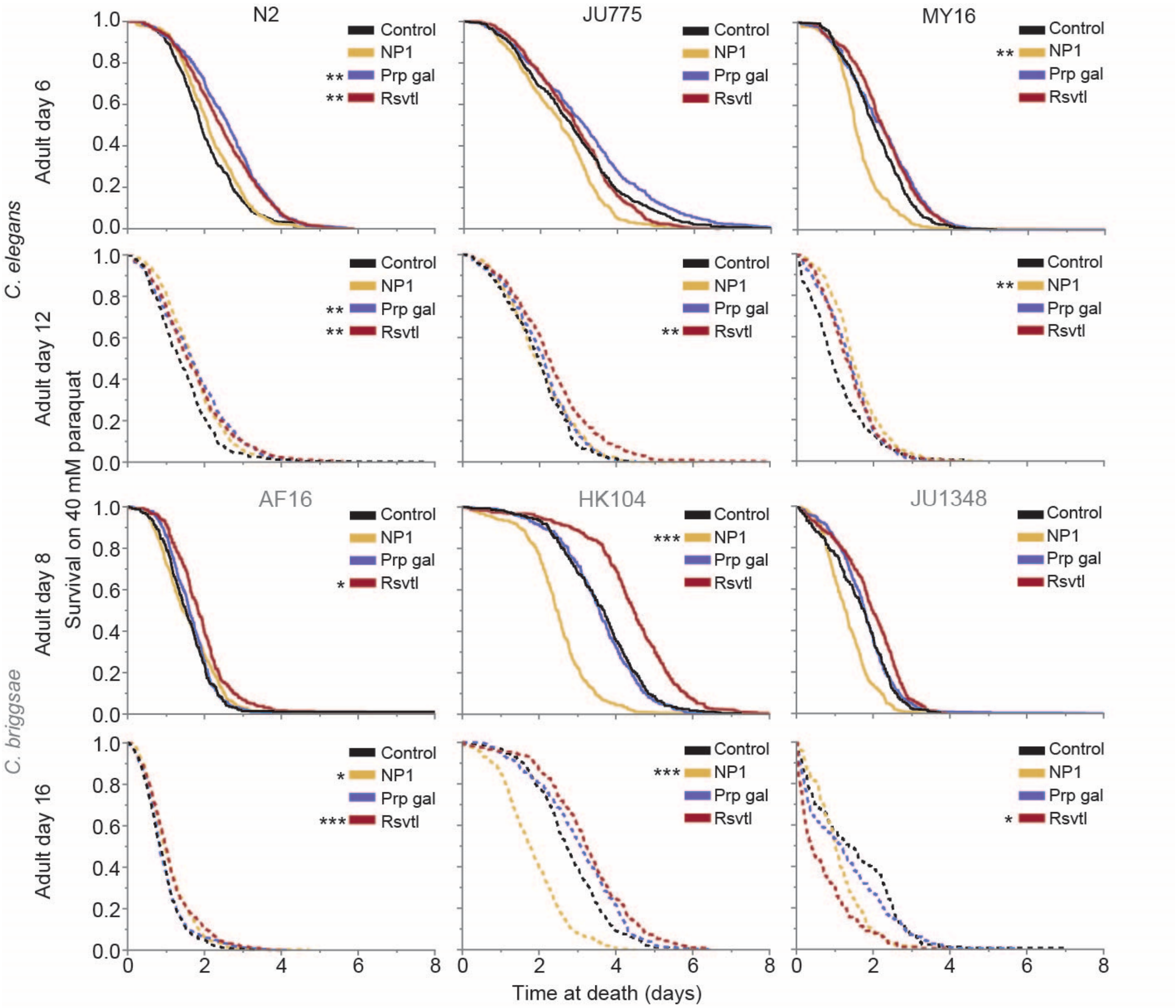


**Supplemental Figure 2. Oxidative stress supplemental**: Kaplan-Meier curves showing the effect of adult exposure to NP1, propyl gallate, or resveratrol on survival under oxidative stress conditions (40 mM paraquat) beginning at day 6 and 12 (*C. elegans*), or day 8 and 16 (*C. briggsae*) of adulthood. Solid lines indicate the younger age, dashed indicate the older age. Each curve represents multiple biological and technical replicates conducted at each of the three CITP testing sites. Asterisks represent *p*-values from the CPH model such that ****$p<0.0001$, ***$p<0.001$, **$p<0.01$, and *$p<0.05$.

### Supplemental Figures

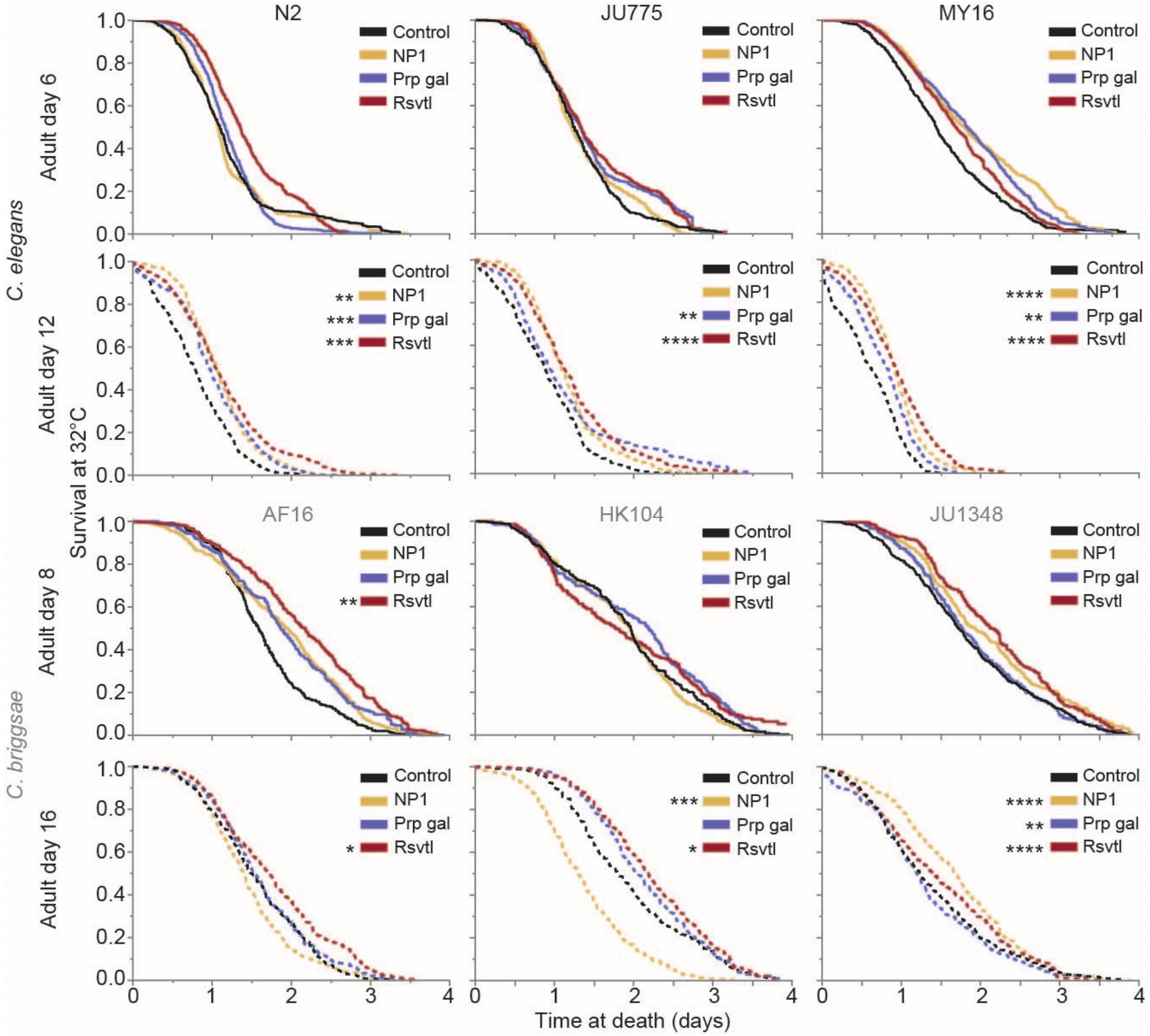


**Supplemental Figure 3. Thermotolerance supplemental**: Kaplan-Meier curves showing the effect of adult exposure to NP1, propyl gallate, or resveratrol on thermotolerance at 32°C beginning on day 6 and 12 (*C. elegans*) or day 8 and 16 (*C. briggsae*) of adulthood. Solid lines indicate the younger age, dashed indicate the older age. Each curve represents multiple biological and technical replicates conducted at each of the three CITP testing sites. Asterisks represent *p*-values from the CPH model such that ****$p<0.0001$, ***$p<0.001$, **$p<0.01$, and *$p<0.05$.

### Supplemental Figures

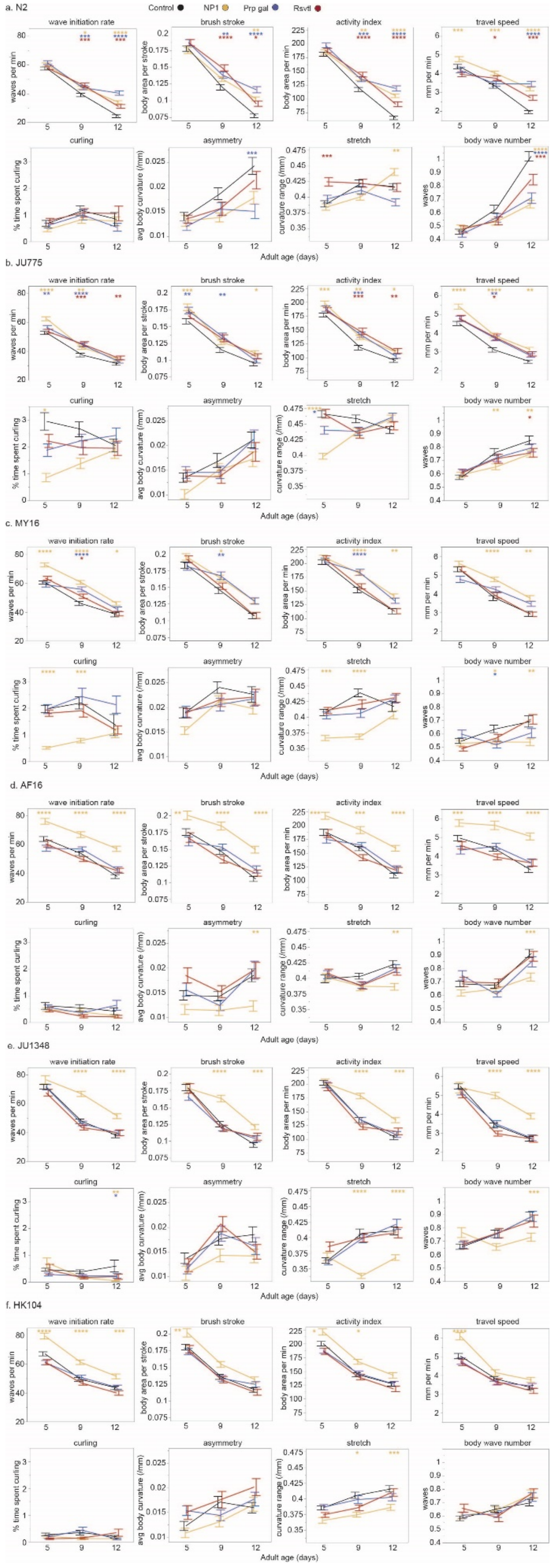

**Supplemental Figure 4. CeleST Panel of Eight Measures:** The effect of adult exposure to NP1, propyl gallate, or resveratrol on eight measures of swimming ability with age. Data is shown for three *C. elegans* strains: (a) N2, (b) JU775, and (c) MY16, and three *C. briggsae* strains: (d) AF16, (e) JU1348, and (f) HK104. Swimming assays were run at early mid-life, mid-life, and late mid-life (days 5, 9, and 12 of adulthood, respectively). The line represents the mean of an individual trial, bars represent the mean +/- the standard error of the mean, and the colors correspond to the compound treatment (black- control, yellow- NP1, purple- propyl gallate, red- resveratrol). Two biological replicates were completed at each of the three CITP testing sites. Asterisks represent *p*-values from the linear mixed model such that ****$p<0.0001$, ***$p<0.001$, **$p<0.01$, and *$p<0.05$.

### Supplemental Figures

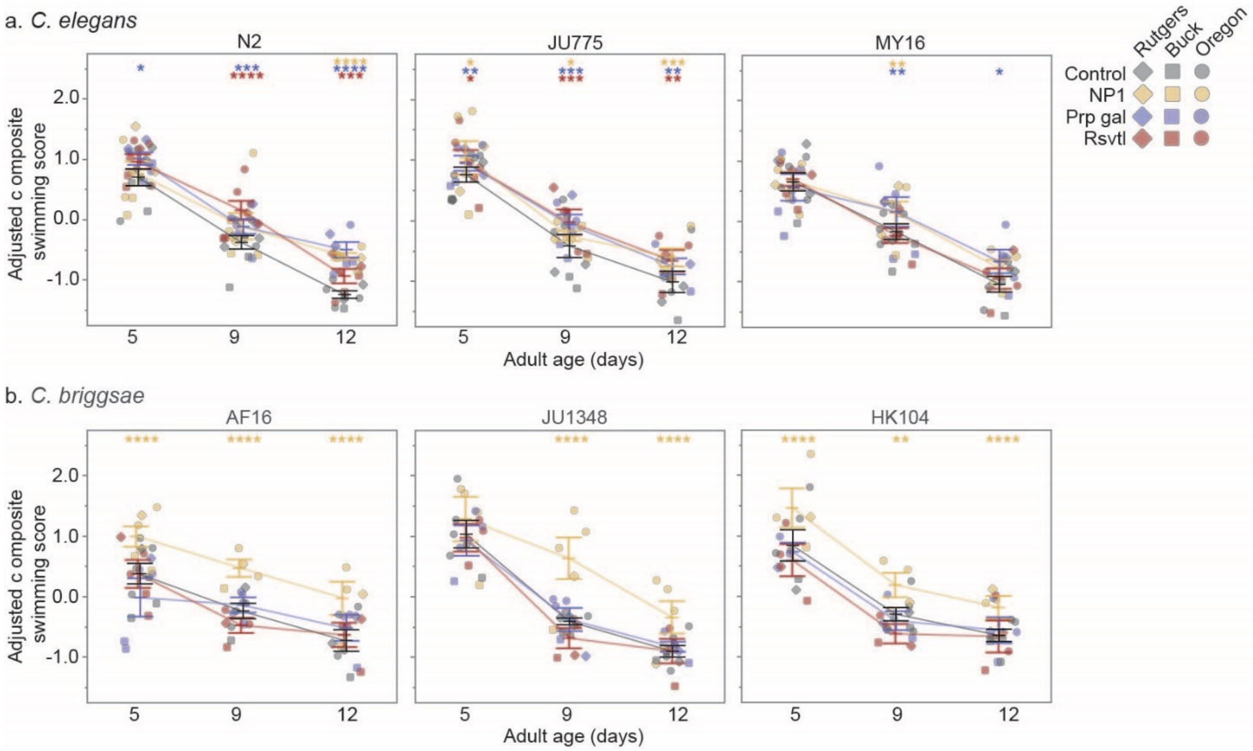


**Supplemental Figure 5. CeleST composite supplemental:** The effect of adult exposure to NP1, propyl gallate, or resveratrol on overall swimming ability with age in three *C. elegans* strains (N2, JU775, MY16), and three *C. briggsae* strains (AF16, JU1348, HK104). Swimming assays were run on days 5, 9, and 12 of adulthood. Points represent the mean of an individual trial, bars represent the mean +/- the standard error of the mean, and the colors correspond to the compound treatment (black – control, yellow – NP1, purple – propyl gallate, red – resveratrol). Adjusted swimming score values were normalized to the strain mean value. Two biological replicates were completed at each of the three CITP testing sites. Asterisks represent *p*-values from the linear mixed model such that ****$p<0.0001$, ***$p<0.001$, **$p<0.01$, and *$p<0.05$.

### Supplemental Figures

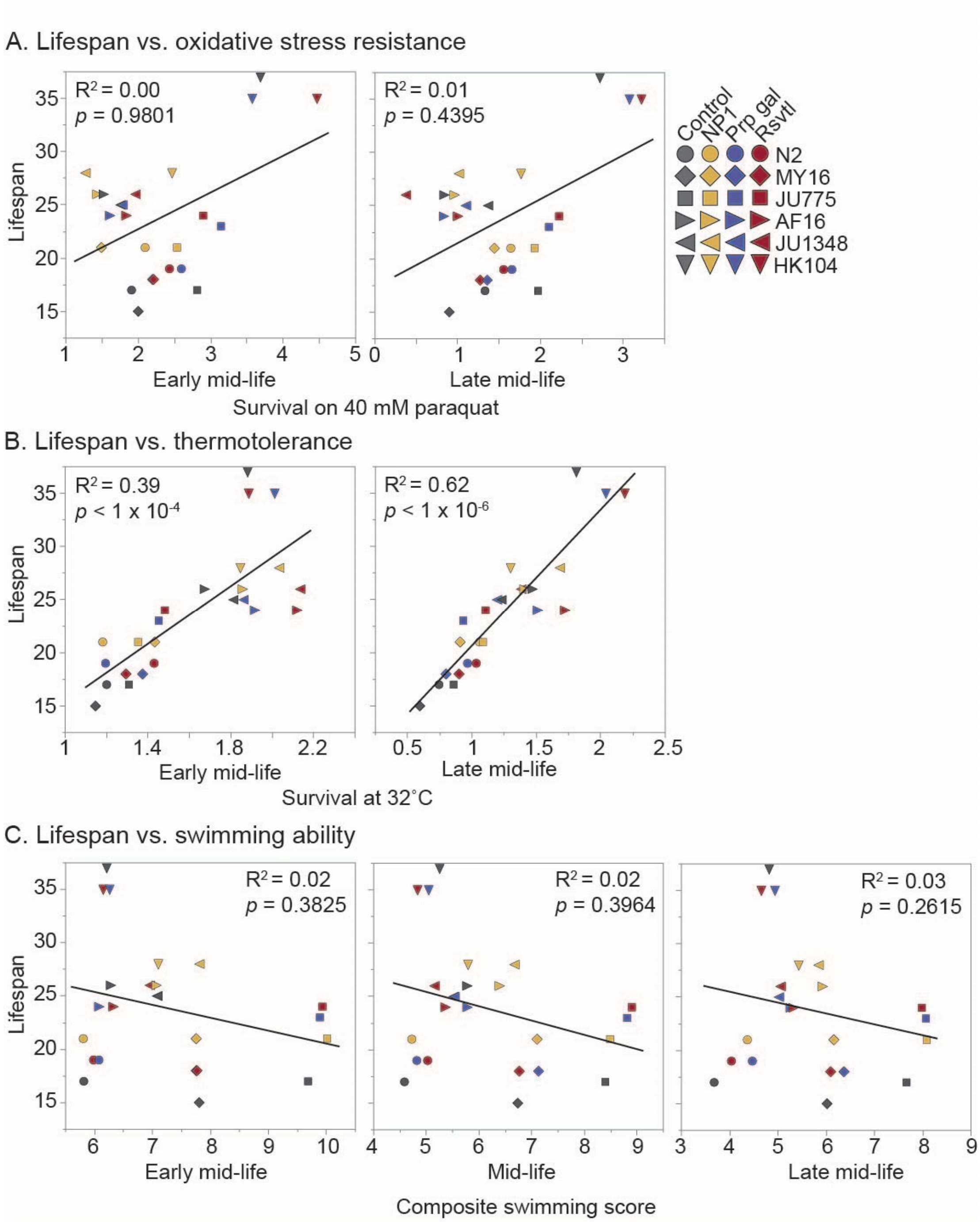


**Supplemental Figure 6. Correlation between lifespan and age-specific health measures of compound intervention:** Health was measured at early and late mid-life for all assays, with an additional measurement at mid-life for swimming ability. Median values were used for composite swimming ability, while Kaplan-Meier medians were used for lifespan, oxidative stress resistance, and thermotolerance. Kendall's tau correlation coefficients were calculated using data from three *C. elegans* strains (N2, MY16, and JU775) and three *C. briggsae* (AF16, JU1348, and HK104) across multiple compound interventions (DMSO controls, NP1, propyl gallate and resveratrol). The only significant correlation with lifespan was for thermotolerance, with $R^2 = 0.39$ for early mid-life, and $R^2 = 0.62$ at late mid-life ($p$<0.0001).
