## Supplemental File 1 for "The coupling between healthspan and lifespan in *Caenorhabditis* depends on complex interactions between compound intervention and genetic background"

### The coupling between healthspan and lifespan in *Caenorhabditis* depends on complex interactions between compound intervention and genetic background

Stephen A. Banse[1&], E. Grace Jackson[1&], Christine A. Sedore[1&], Brian Onken[2], David Hall[3], Anna Coleman-Hulbert[1], Phu Huynh[2], Theo Garrett[3], Erik Johnson[1], Girish Harinath[2], Delaney Inman[3], Suzhen Guo[2], Mackenzie Morshead[3], Jian Xue[2], Ron Falkowski[2], Esteban Chen[2], Christopher Herrera[2], Allie Kirsch[1], Viviana I. Perez[4], Max Guo[4], Gordon J. Lithgow[3*], Monica Driscoll[2*], Patrick C. Phillips[1*]

#### *Generation of a Composite CeleST score*

Locomotion is an important health measure for animals ranging from flies to humans[1–3]. For *Caenorhabditis elegans*, plate-based locomotion measures, typical mean and maximum velocity, are commonly used health measures[4,5]. The *Caenorhabditis* Intervention Testing Program (CITP)[6] adopted the CeleST platform[7–9], which acquires eight measures of swimming ability, as a more informational deep locomotion assay (see Supplemental Figure 4). While the eight measures provide a broad range of information on the swimming of each of the strains, it was not clear which measures best capture the decline of locomotion with age across our genetic diversity panel[10]. We therefore combined the information from all measures for each strain into a single multivariate composite measure by developing an analysis program pipeline that creates a single composite swimming score from eight original swim test measurements.

The data used to generate the composite score were generated by processing 30 second videos with the published CeleST software[7,8]. The software provides eight measurements of mobility, like travel speed, asymmetry, curling, and body waves initiation, for individual worms. We collected the data at three days in adulthood (5, 9, and 12). To generate a composite score, we first analyzed the eight original measurements in each record using a linear discriminant analysis (LDA) approach using the eight original measurements in each record as the predictor variables and the age-dependent decline as the primary discriminator. By projecting the eight predictor variables onto a single axis, LDA creates the first linear discriminant function which maximizes the differences between ages across all measurements, while reducing the dimensionality from eight to one (see visual example).

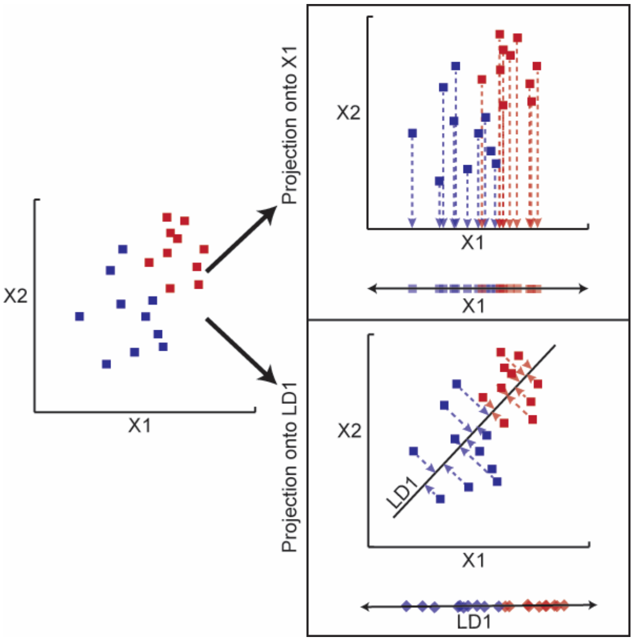


**Visual example of LDA dimensional reduction:** In this example there are two predictor variables (X1 and X2) and two levels to the group (red and blue). All of the information captured is in the leftmost plot of variable X1 x X2. To reduce the number of dimensions down to one, we would project that plot onto a single axis. If projecting to one of the original axes, e.g., the X1 axis presented in the example, all information from the second variable is lost and little benefit in distinguishing between levels is created. By creating a new axis to project onto, LD1 in the example shown, we preserve nearly all of the information from both variables while still reducing the dimension to one, and that new first linear discriminate function has a coefficient for each of the original predictor variables.

**Supplemental File 1**

The first linear discriminant function minimizes within-group (age in our case) variance, maximizes between-group (age) variance, and maximizes the separability of the means of the groups (ages), while having a coefficient for each of the original eight predictor variables[11]. While it cannot capture *all* the information provided by the original measurements, it does maximize informational capture. Because this approach maximizes the differences between group levels, linear discriminant analysis is often used to predict group membership after a training dataset is used. Generating the coefficients for each of the original eight predictor variables in a strain specific manner enabled us to avoid arbitrarily determining how any of the eight measures vary between ages, or if they matter at all in differentiating ages, while avoiding the assumptions that the strains swim and age the same.

To implement the LDA approach for analyzing compound interventions, LDA is first performed on the control dataset, independently for each strain, to retrieve the coefficients from the first linear discriminant function. The strain-specific coefficients are then used as weighted loadings which are multiplied by the corresponding measurement and divided by the control dataset's measurement standard deviation. This is done for every worm record in the strain dataset, including compound and control records. Those eight weighted measurements are summed up within each record to create the composite swimming score. This composite score can then be used to compare compound interventions and ages within each strain without arbitrarily choosing measurements for comparison. For statistical comparison, we used models with compound treatment and age as an interaction for input into Type-III Analysis of Variance tests for the effect of a factor, given that other factors are present. A significant age by compound interaction is of the most interest, as it implies that the impact size of a compound varies by age. The scripts used for generating the composite scores and analysis, and additional relevant information, have been made available online[12].

*CeleST Composite Score Example*

Because each record has eight measurements and belongs to a particular strain, compound, and age, we were presented with question: what do these measures mean in terms of health? Our solution is to combine the information provided by these measurements while considering (i) the relevance of each measure, (ii) the different units and scales, and (iii) how not all measures are independent. By combining the information into a single value using the LDA approach described above, we avoided arbitrary choices. Below we show an example for the processing of sample data.

The example below is from the N2 portion of the dataset included in this manuscript. In this case, Activity Index is the most important measure across the board when it comes to age and will have a greater impact on the final score. Curling and Travel Speed are nearly irrelevant and will not impact the score much. These loadings vary by strain and can be negative or positive. The loadings were negated from the initial output, so the direction of the score goes down over time for any particular strain-age-compound combination. LDA doesn't see each age as older/younger, just as distinct groups, so negating those loadings is the same as flipping the order of the age levels.

$$= \sum_{n=1}^{8} Measurement_n \frac{Loading_n}{DMSO\ SD_n}$$

| N2 | Wave | Brush | Activity | Travel | Curling | Asymmetry | Stretch | Body |
|---|---|---|---|---|---|---|---|---|
| Loading | 0.355 | 0.235 | 1.154 | -0.002 | -0.046 | 0.169 | 0.327 | 0.336 |
| DMSO SD | 23.912 | 0.077 | 78.208 | 2.172 | 3.321 | 0.023 | 0.096 | 0.557 |

**Supplemental File 1**

Now when we look at the composite swimming score for each strain-age-compound combination, much of the relevant information from the eight original measurements is captured, and we can focus our analyses on that single new measurement. Similar to our work with survival/lifespan data[6], we create two mixed effects general linear models (GLM) for each strain:

CompositeSwimmingScore ~ Compound * Age + (1|Lab/Tech/Date/Video)

CompositeSwimmingScore ~ Compound_Age + (1|Lab/Tech/Date/Video)

The score is the response in the same way that death age/time is the response with lifespan data. The fixed effects term has an interaction, and the random effects are nested instead of crossed. This means we break the random effects levels into a tree – with an experiment date at one lab is unique from the same date at a different lab.

To look at the significance of an interaction between compound and age, a Type-III ANOVA is performed on the first GLM. This type of ANOVA tests for the effect of a factor, given that the other factors are present. This approach is best when a significant interaction effect is present, as the effect of that interaction is tested after considering the individual factors. The presence of a significant interaction effects suggests the score is telling us something about healthspan, not just lifespan. A single test is run for each strain. Any breakdown of the sources of variance uses a similar GLM with the interaction term.

To compare between compound and ages, the second GLM is used. This GLM is similar to the first, but instead of having an interaction term, the response variable combines compound and age. That way the control and the compound can be compared at particular ages. These pairwise comparisons are for the differences between the means of the Composite Swimming Score. All analyses are done within a particular strain.
